# Establishment of a hypoxia ischemia reperfusion brain damage model in neonatal rats

**DOI:** 10.1101/2022.01.10.475606

**Authors:** Xiaoqin Fu, Tianlei Zhang, Wei Lin, Mengdie Jiao, Zhiwei Zhang, Jiayi Geng, Qing Wei, Ailin Qi, Lin Kexin, Yi hui Zheng, Mingchu Fang, Peijun Li, Zhenlang Lin

## Abstract

**Objective:** Rice-Vannucci model has been widely used as HIE (Hypoxic ischemic encephalopathy) animal model in the past forty years, but it does not mimic reperfusion injury that occurs during HIE. The aim of the present study was to establish a new neonatal rat model by simulating hypoxia ischemia reperfusion brain damage (HIRBD) through “common carotid artery (CCA) muscle bridge”.

**Methods:** Sixty 7-day-old male Sprague-Dawley rats were randomly assigned to group A (HIRBD groups, n=36), group B (Rice-Vannucci group, n=12), and group C (sham-operated group, n=12). Rats in group A were assigned to 3 subgroups (A1-A3, 12 animals/subgroup). Dynamic changes in cerebral blood flow (CBF) were evaluated by the laser speckle imaging system. The status of the CCA was observed under a stereomicroscope. Changes in body weight, gross morphology as well as pathological sections of brain tissue were examined to evaluate the feasibility of the model.

**Results:** The results indicated that CCA muscle bridge successfully blocked the CBF. CBF was restored after removal of the CCA muscle bridge in HIRBD groups. The CCA was in good condition after removing the muscle bridge, and blood supply was not affected. Changes in body weight, gross morphology and pathological sections of brain tissue indicated that ischemia reperfusion induced by the CCA muscle bridge method caused varying degrees of brain damage.

**Conclusion:** CCA muscle bridge method is effective for establishing a reliable, stable, and reproducible neonatal rat model for study of HIRBD.

## Introduction

Hypoxic ischemic encephalopathy (HIE) is a devastating disorder caused by perinatal asphyxia. HIE leads to hypoxic ischemic brain damage (HIBD) in fetuses and neonates. Hypoxia is the main cause of pathophysiological changes in HIE, including reduction in cerebral blood flow (CBF), decrease in cerebral vascular autonomic function, failure of cell energy metabolism, necrosis and apoptosis caused by free radical damage, intracellular calcium overload and production of excitatory amino acid and cytokines. These factors can lead to structural and functional damage in the brain^[1,2]^. Studies report that ischemia reperfusion (IR) induced brain injury is an important factor in induction of neonatal neurological sequelae^[3]^. IR involves multiple cellular mechanisms including oxidative stress, ferroptosis, neuron protein synthesis inhibition, mitochondrial dysfunction, and inflammatory response^[4–8]^. Studies exploring ischemia reperfusion induced brain injury, secondary to hypoxic ischemia in the pathology of HIE, will offers a potential therapeutic target for management of HIBD^[3,9]^.It is thus imperative to explore the pathogenesis and treatment of HIRBD.

Animal models play significant roles in exploring pathogenesis and treatment of HIE. Rice and Vannucci’s landmark study^[10,11]^ reported a classic neonatal rat model of HIBD, which was simple to establish. However, the model has some drawbacks such as: it does not achieve ideal cerebral hemisphere ischemia owing to existence of Circle of Willis. In addition, Rice-Vannucci model does not cause reperfusion injury since it involves permanent ligation. Therefore, we designed a new neonatal rat model that simulates HIRBD through “common carotid artery (CCA) muscle bridge”.

## Materials and Methods

### Instruments and reagents

Instruments used in the current study included binocular stereomicroscope (XTZ-D, Cai Kang Optical Instrument Co. Ltd. Shanghai, China), hypoxia chamber (Hangzhou Aipu Instrument and Equipment Co. Ltd. Zhejiang, China), thermostatic incubator (Thermo Forma, USA), laser speckle imaging system (RFLSI III, RWD Life Science, Shenzhen, China) and paraffin slicer (Leica, Germany). Reagents used the present study included Nissl staining solution (Beyotime Biotechnology Co. Ltd. Shanghai, China), Hematoxylin-Eosin staining solution (Beyotime Biotechnology Co. Ltd. Shanghai, China), rabbit anti-rat glial fibrillary acidic protein (GFAP) polyclonal antibody (ProteinTech, Wuhan, China) and FITC - goat anti-rabbit IgG (Nanjing Kaiji Biological Technology Development Co. Ltd. Nanjing, China).

### Animal studies

All animal experiments were performed in strict accordance with the guidelines of the Care and Use of Laboratory Animals and were approved by the Animal Experimentation Ethics Committee of Wenzhou Medical University. Sprague-Dawley (SD) rats, weighing 200-250g, were obtained from Weitong Lihua Experimental Animal Center (Beijing, China) and were housed under specific-pathogen-free (SPF) conditions. Adult rats were allowed to freely mate to produce offspring for subsequent studies. A total of 60 male rats at postnatal day 7 (P7) were used in the study. The weights of the rats ranged from 13-18g (no statistically significant differences among the groups).

Animals were anaesthetized by 3%-4% isoflurane inhalation (maintained with 1%-2% isoflurane) before performing surgeries. Animals were allowed to recover from anesthesia for 10 minutes (rats generally recover from anesthesia within 5 minutes) after surgery. Pups were returned to their dam and remained with their mother after experiments.

### HE staining

Seventy-two hours after HIBD modeling, the rats in both groups were deeply anesthetized with isoflurane, and the hearts were perfused with 20 ml of saline, followed by 20 ml of 4% PFA. Immediately after perfusion, the heads of the rats were cut off, the scalp and skull were cut, and the brain tissue were preserved in 4% PFA solution at 4°C, dehydrated and embedded in paraffin. Further, 4μm thick coronal sections were made from the tissues. The sections were dewaxed, stained with hematoxylin, differentiation solution and eosin, then permeabilized in xylene. Sections were then sealed using neutral balsam, and the morphology of brain tissues was viewed under a microscope.

### Nissl staining

Tissue sections were dewaxed, stained using Cresyl violet acetate and differentiation solution, then permeabilized in xylene. The sections were then sealed with neutral balsam, and the morphology of brain tissues was viewed under a microscope.

### Immunofluorescence

Sections were dewaxed then heated in citrate buffer for antigen retrieval. Sections were then permeabilized in Triton x-100, rinsed with PBS and blocked with 10% goat serum.

Sections were incubated with rabbit anti-rat GFAP polyclonal antibody (1:200) overnight at 4°C. The sections were then rinsed with PBS again and incubated with FITC-goat anti-rabbit IgG for 90 minutes at 37°C. Sections were sealed after rinsing with PBS and observed under a laser microscope.

### HIBD rat model

A Rice-Vannucci HIBD model was established as described previously. The left CCA were exposed under stereomicroscope. The proximal and telecentric ends of the left CCA were ligated with 8-0 sutures, and carefully cutting the artery between the ligation knots. Rats were allowed to rest in an incubator at 37°C for 10 minutes after surgery. Further, rats were subjected to hypoxia for 120 minutes in a hypoxia chamber (8% oxygen and 92% nitrogen). The left common carotid artery of each pup in the sham surgery group was only identified, exposed and isolated, and the pups were not subjected to hypoxia.

### HIRBD animal grouping

Neonatal rats from each litter were randomly assigned to three groups: group A (HIRBD group, subdivided into A1, A2 and A3 subgroups), group B (Rice-Vannucci HIBD group), and group C (sham-operated group), with 12 rats in each group. Details on the grouping are presented in Table 1.

**Table 1.** Time of hypoxia and ischemia induction in each group

| Group |  | Surgical operations | CCA | Ischemic time (minute) | Hypoxic time (minute) |
| --- | --- | --- | --- | --- | --- |
| A (HIRBD group) | A1 | Left CCA ligated and severed; right CCA-muscle-bridge | Left | Permanent ischemia | 30 |
|  |  |  | Right | 90 |  |
|  | A2 | Bilateral CCA-muscle-bridge | Left | 90 | 30 |
|  |  |  | Right | 90 |  |
|  | A3 | Right CCA-muscle-bridge | Left | No processing | 120 |
|  |  |  | Right | 150 |  |
| B (HIBD group) |  | Rice-Vannucci HIBD model | Left | Permanent ischemia | 120 |
|  |  |  | Right | No processing |  |
| C (Sham group) |  | Sham-operated | Left | No processing | N/A |
|  |  |  | Right | No processing |  |

### HIRBD rat model

#### 1) CCA muscle bridge (right CCA muscle bridge is used as an example)

Rats were fixed on a sterile animal operating table after anaesthetization. An incision was made at the midline after disinfection of the neck skin. Bilateral CCA, bilateral mastoid muscle (MM) and stemohyoideus muscle (SHM) were exposed under a stereomicroscope. The SHM was then bisected from the middle into two fascicles (left and right fascicle). One end (defined as A, Figure 1h) of a 5cm 8-0 suture line was orderly passed through the underside of the right MM, the right CCA and the right fascicle of SHM. The other end of the suture line crossed over the right MM then passed through the underside of the right CCA. The two ends of the suture line were tied together above the right fascicle of the SHM, such that the right MM and the right fascicle of the SHM were joined together to form the first “muscle bridge” which shored up and compressed the right CCA. The second “muscle bridge” was established 3mm from the telecentric end of the first CCA “muscle bridge”. A 5cm 8-0 suture line (defined as B, Figure 1h) was passed over the right CCA, then crossed below the right MM and the right fascicle of SHM. The other end of the suture line orderly passed over the right MM, the right CCA and the right MM. The two ends of the suture line were joined above the right CCA. The two “muscle bridge”, the right MM and the right fascicle of SHM were tied together to “clamp” the CCA (Figure 1f).

**Figure 1.**
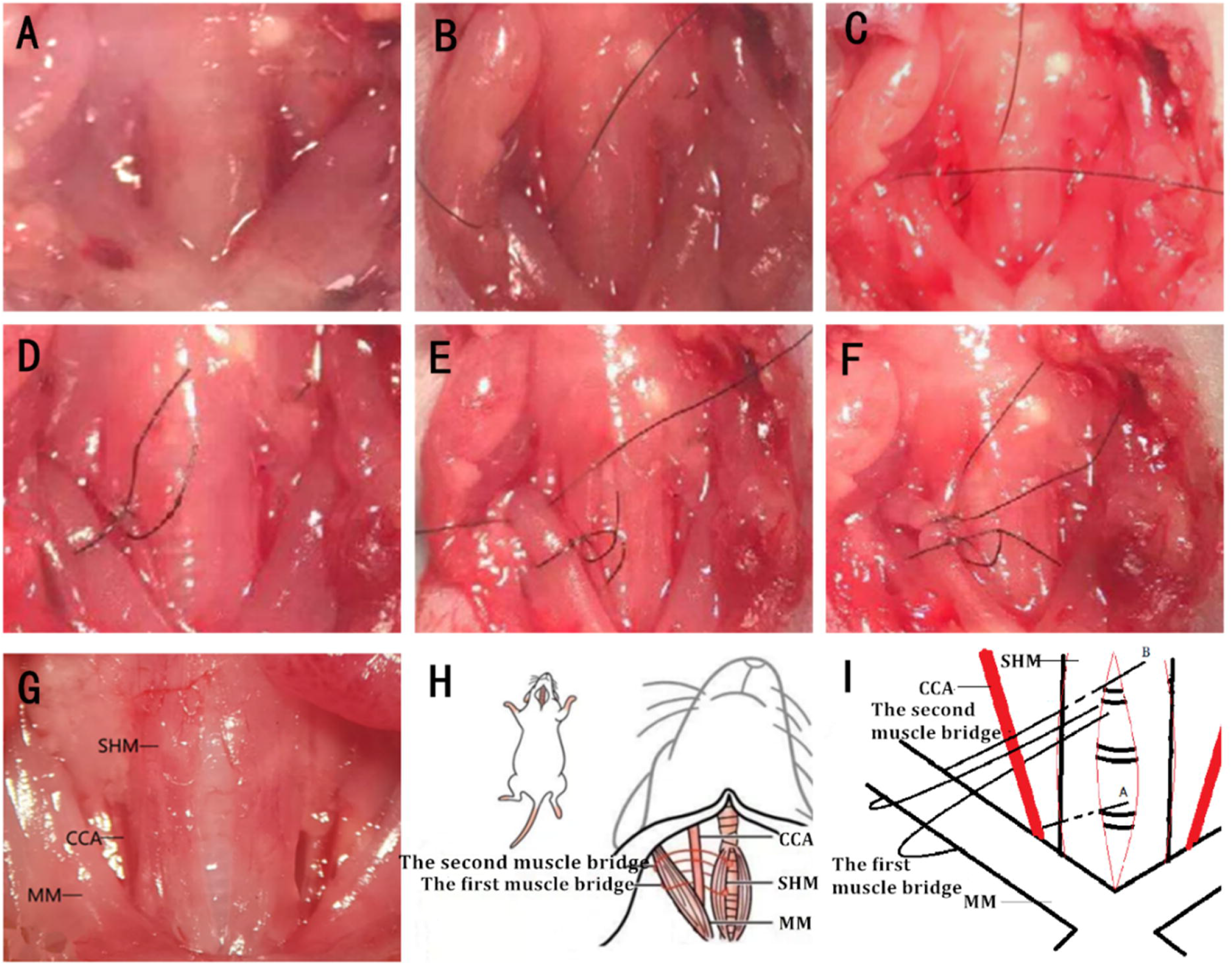
Illustration of establishment of CCA muscle bridge (A).Bilateral MM and SHM in front of the trachea. (B). The first suture line passed in turn through the underside of the right MM, the right CCA and the right fascicle of SHM. (C). The other end of the first suture line was passed over the right MM then passed through the underside of the right CCA. (D). After driving the line through the muscle, the two ends of the first suture line were tied together above the right fascicle of the SHM to form the first muscle bridge. (E). The second suture line was passed above the right CCA, then passed through the underside of the right MM and the right fascicle of SHM. (F). The second suture line were was tied at its two ends above the right CCA to form the second muscle bridge. (G). Morphology of MM, CCA and SHM incised from center. (H) Illustration of the process of establishing CCA muscle bridge. (I) Effect diagram of the first and the second CCA muscle bridge.

Muscle bridge relief: The knots of bridges were cut with microsurgical scissors after removing the suture of the original incision to relieve the compression on the CCA during the predetermined ischemic time. The incision was sutured after restoring blood flow and rats were allowed a 10-minute-break to recover from anesthesia.

#### 2) Time of hypoxia and ischemia induction

Hypoxic exposure was initiated when oxygen concentration reached 8% to the point when experimental animals were removed from the hypoxia chamber. Hypoxic exposure time for group A1 and A2 was 30 minutes to avoid severe damage caused by bilateral CCA ischemia. Hypoxic time for group A3 and B was 120 minutes (Table 1). Ischemia timing of group A1-A3 started from the time the surgery was completed and ended at the time the muscle bridge was relieved. Ischemic time for A1 and A2 was set as 90 minutes, and that for A3 as 150 minutes.

### Evaluation methods for HIRBD

The models were evaluated according to the hemodynamics (by the laser speckle imaging system), the state of the CCA on the third day after modeling (by the stereomicroscope), body weight changes, gross morphology and analysis of pathological sections of brain tissue (HE staining, Nissl staining and GFAP immunofluorescence).

#### 1) Laser speckle imaging technique (LSI)

LSI can be used for detection of the perfusion unit (PU) of cerebral blood flow (CBF) below the skull of neonatal rats. The left and right CBF can be recorded and compared when establishing and relieving the muscle bridge. Rats (group A1, A2, A3, B, C) were anesthetized after induction of hypoxia. The skull was exposed by making an arc-shaped incision in the scalp. The CBF of the left and right hemispheres was then determined under a 785nm wavelength laser. Data were recorded as A1, A2, A3, B and C. The scalp was sutured after surgery and rats were placed back with their mother after waking up from anesthesia. CBF for rats in CCA muscle bridge surgery groups were recorded as A1-1, A1-2, A2-1, A2-2, A3-1 and A3-2, after ischemic treatments and the muscle bridge was removed as described in Table 2. Data were analyzed using LWCI software (RFLSI III, RWD Life Science, Shenzhen, China).

**Table 2.** LSI test time for each group

|  | Left CCA | Right CCA | Test time |
| --- | --- | --- | --- |
| A1-1 | Ligated and cut | muscle bridge established | at the end of hypoxia treatment |
| A1-2 | Ligated and cut | muscle bridge removed | after the remove of the muscle bridge |
| A2-1 | Muscle bridge established | muscle bridge established | at the end of hypoxia treatment |
| A2-2 | muscle bridge removed | muscle bridge removed | after the remove of both muscle bridges |
| A3-1 | No operation | muscle bridge established | at the end of hypoxia treatment |
| A3-2 | No operation | muscle bridge removed | after the remove of the muscle bridge |
| B | Ligated and cut | No operation | at the end of hypoxia treatment |
| C | sham surgery | No operation | After sham operation |

#### 2) Status of CCA

The status of CCA was determined on the third day after modeling. Analysis was conducted to explore whether the bilateral CCA was in good pulsation or not using a binocular stereomicroscope. Further examination was conducted to explore whether muscle bridges affected the fluidity of blood flow of CCA by clipping CCA. The telecentric end of muscle bridge was ligated (the muscle bridge had been removed, thus the ligation position was selected far away from the distal end of the CCA of the original position of the muscle bridge). The CCA was then at the proximal end close to the ligation to observe the spillover of arterial blood.

#### 3) Body weight changes

Body weight of the rats was recorded daily. Body weight was compared for 3 days after modeling of each group.

#### 4) Analysis gross morphology and pathological sections of brain tissue

The degree of cerebral injury was evaluated by analysis of gross morphology, HE staining, Nissl staining and GFAP fluorescence staining of brain tissue sections. Brain sections were classified into four grades: mild, moderate, severe, and very severe (Table 3). Brain tissue was classified at a particular grade with the corresponding score, if it met two of the three items in the scoring table, then the mean score of each group was finally calculated.

**Table 3.** Degree and score of cerebral injury

| Score | Degree | Morphology | HE staining/Nissl staining | GFAP immunofluorescence |
| --- | --- | --- | --- | --- |
| 0 | Normal | Liquefaction (-)<br>atrophy (-)<br>edema (-) | Neurons show a clear hierarchy, with the integrity of structure as well as normal number and morphology. The cell membranes are intact. The nuclei are centered, round and contain deeply stained nucleus; Nissl bodies are densely and deeply stained. | Few GFAP positive cells are observed in the Corpus callosum |
| 1 | Mild | Liquefaction (-)<br>atrophy (-)<br>edema (-) | Same as above | Some GFAP positive cells are observed in the Corpus callosum |
| 2 | Moderate | Liquefaction (-)<br>atrophy (+)<br>or edema (+) | Mild atrophy and mild reduction of the number of neurons in cerebral cortex and hippocampus. A small number of neurons show morphological changes. Necrotic neurons appeared locally, and Nissl body staining become lighter. | Same as above |
| 3 | Severe | atrophy (++)<br>or edema (++) | Severe atrophy and reduction of neurons observed, as well as small patches of liquefactive necrosis. The cell bodies of neurons are swollen. Local lesions exhibit pyknotic nuclei of necrotic neurons, and detritus appears; Nissl body staining is weak and sparse. Large patches of Nissl bodies are vanished. | Same as above |
| 4 | Very Severe | liquefactive necrosis (+) | Large patches of liquefactive necrosis are observed in the sections. | GFAP-positive cells are scattered in the liquefied area |

### Gross morphology analysis

Rats were cardiac perfused with 20ml PBS solution after anesthesia. Brains were then harvested from the skull, followed by imaging to evaluate gross morphology. Brain tissues were then preserved in PFA solution at 4°C.

### HE staining

Brain tissues were preserved in 4% PFA solution at 4°C then dehydrated and embedded in paraffin. Further, brain tissues were cut into 4μm thick coronal sections. The sections were dewaxed, stained with hematoxylin, differentiation solution and eosin, then permeabilized in xylene. Sections were subsequently sealed with neutral balsam, and the histopathology of brain tissues observed under a microscope.

### Nissl staining

Sections were dewaxed, stained using Cresyl violet acetate and differentiation solution, then permeabilized in xylene. The sections were then sealed with neutral balsam, and histopathology of brain tissues explored under a microscope.

### Immunofluorescence analysis

Sections were dewaxed and heated in citrate buffer for antigen retrieval. Further, sections were permeabilized in Triton x-100, rinsed with PBS and blocked with 10% goat serum. Sections were incubated with rabbit anti-rat GFAP polyclonal antibody (1:200) at 4°C overnight. Sections were then rinsed with PBS and incubated with the FITC-goat anti-rabbit IgG for 90 minutes at 37°C. Sections were rinsed with PBS and sealed, then observed under a laser microscope.

### Cell count of hippocampus

Cells in the symmetrical hippocampus of HE-stained sections were counted. Six fields (40 ×) of the sections were selected for cell counting to compare the degree to damage of the hippocampus.

### Statistical analysis

Quantitative data were obtained from at least three independent experiments. Comparison between two groups was performed using the Student’s t-test. Differences between more than two groups were compared through analysis of variance (ANOVA). All statistical analyses were performed using IBM SPSS Statistics 21.0.0.0 (International Business Machines Corporation., USA). The level of statistical significance was set at p<0.05.

## Results

### **Mortality** the rate of HIRBD was not significantly different compared with that of HIBD

The A1(2/12) showed the highest mortality rate. Mortality rates were all 1/12 in group A2, A3 and B (Table 4). The left brain of group A1 was subjected to permanent ischemia, whereas the right brain was subjected to transient ischemia treatment, thus the injury in group A1 was the most severe and mortality rate was highest in group A1 compared with that of the other groups. Group A3 and B were subjected to the same unilateral ischemia thus had similar mortality rates. These results indicate that the mortality rate of HIRBD was not significantly different compared with that of HIBD.

**Table 4.** Mortality of neonatal rats within 3 days

| Group | Quantity | The number of deaths | Mortality (%) |
| --- | --- | --- | --- |
| A1 | 12 | 2 | 16.67 |
| A2 | 12 | 1 | 8.33 |
| A3 | 12 | 1 | 8.33 |
| B | 12 | 1 | 8.33 |
| C | 12 | 0 | 0 |

#### LSI results

CCA blocks cerebral blood flow, and its effect is similar to that of ligation. Notably, cerebral blood flow is restored after relieving the CCA muscle bridge.

The status of cerebral ischemia and reperfusion in different models was assessed by examination of physical images, speckle images and pseudo color images obtained from LSI (Figure 2). Analysis of pseudo color images of the blood flow before (A*-1) and after (A*-2) removal of CCA muscle bridge in A1-A3 groups showed that cerebral ischemia was successfully established or relieved by the setup or through removal of CCA muscle bridge, respectively. Further quantitative analysis of LSI data showed no statistically significant differences in CBF between the left and right hemispheres in A1-1 group (*P*=0.259). However, the CBF of the left and right hemispheres were significantly different after the right CCA muscle bridge was removed in A1-2 group (*P* < 0.001). The results showed no statistically significant differences in CBF between the left and right hemispheres with bilateral CCA muscle bridge in A2-1 group (*P* = 0.111) and there were also no statistically significant differences in CBF between the left and right hemispheres after removing bilateral CCA muscle bridge in A2-2 group(*P* = 0.088). Analysis of group A3 showed significant differences in CBF between the left and right hemispheres before removal of CCA muscle bridge (*P*=0.001). However, no significant differences were observed after removal of the right CCA muscle bridge in A3-2 (*P*=0.314). The findings for group B showed significant differences in CBF between the left and right hemispheres (*P*<0.001). Analysis showed no statistically significant differences in CBF between the left and right hemispheres for group C (*P*=0.072, Figure 2).

**Figure 2.**
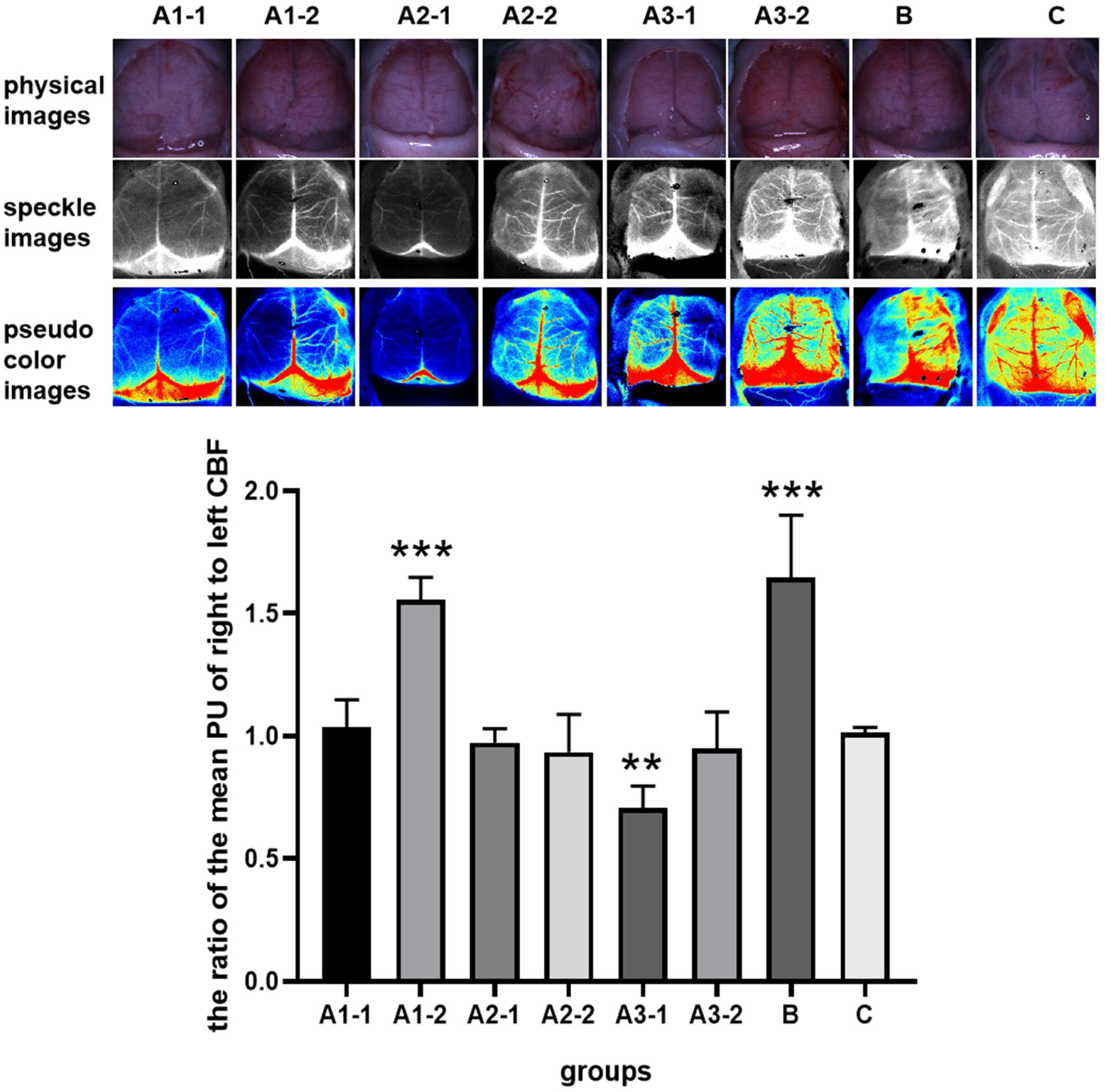
Transient cerebral ischemia was successfully established using CCA muscle bridge. Physical images, speckle images and pseudo-color images obtained from LSI in different groups. A1-1: bilateral cerebral ischemia induced by establishment of right CCA muscle bridge and damage of left CCA. A1-2: The right hemisphere regained blood flow after removal of the right CCA muscle bridge. A2-1: Bilateral cerebral ischemia induced by establishment of bilateral CCA muscle bridge. A2-2: Bilateral hemisphere regained blood flow after removal of the bilateral CCA muscle bridge. A3-1: Right cerebral ischemia induced by establishment of right CCA muscle bridge. A3-2: The right hemisphere regained blood flow after removal of the right CCA muscle bridge. B: Left cerebral ischemia induced by damage of left CCA. C: Results of the sham group. The bar indicates the ratio of the mean PU of right to left cerebral blood flow for each group (* P < 0.05,* * P < 0.01 and * * * P < 0.001).

### Status of CCA after establishment of muscle bridge

Light stenosis of the right CCA was observed in one animal in group A2 and two animals in group A3. Blood spurted from the CCA in all groups after cutting the CCA (Table 5).

**Table 5.** State of the CCA on the third day after modeling

|  | The right CCA<br>of A1 | The left CCA of<br>A2 | The right CCA<br>of A2 | The right CCA<br>of A3 |
| --- | --- | --- | --- | --- |
| Quantity | 10 | 11 | 11 | 11 |
| CCA stenosis | 0 | 0 | 1 | 2 |
| Blood spurted from the CCA | 10 | 11 | 11 | 11 |

### Bodyweight changes

The body weight of each group was recorded 3 days after operation. The findings showed that the body weights of rats in group A1 and A2 significantly decreased after operation. Group A3 and B showed varying degrees of postoperative weight gain, with the most significant increase observed in group C. Analysis showed that the body weights in each group after surgery were significantly different compared with the weight before surgery (P <0.001, Figure 3).

**Figure 3.**
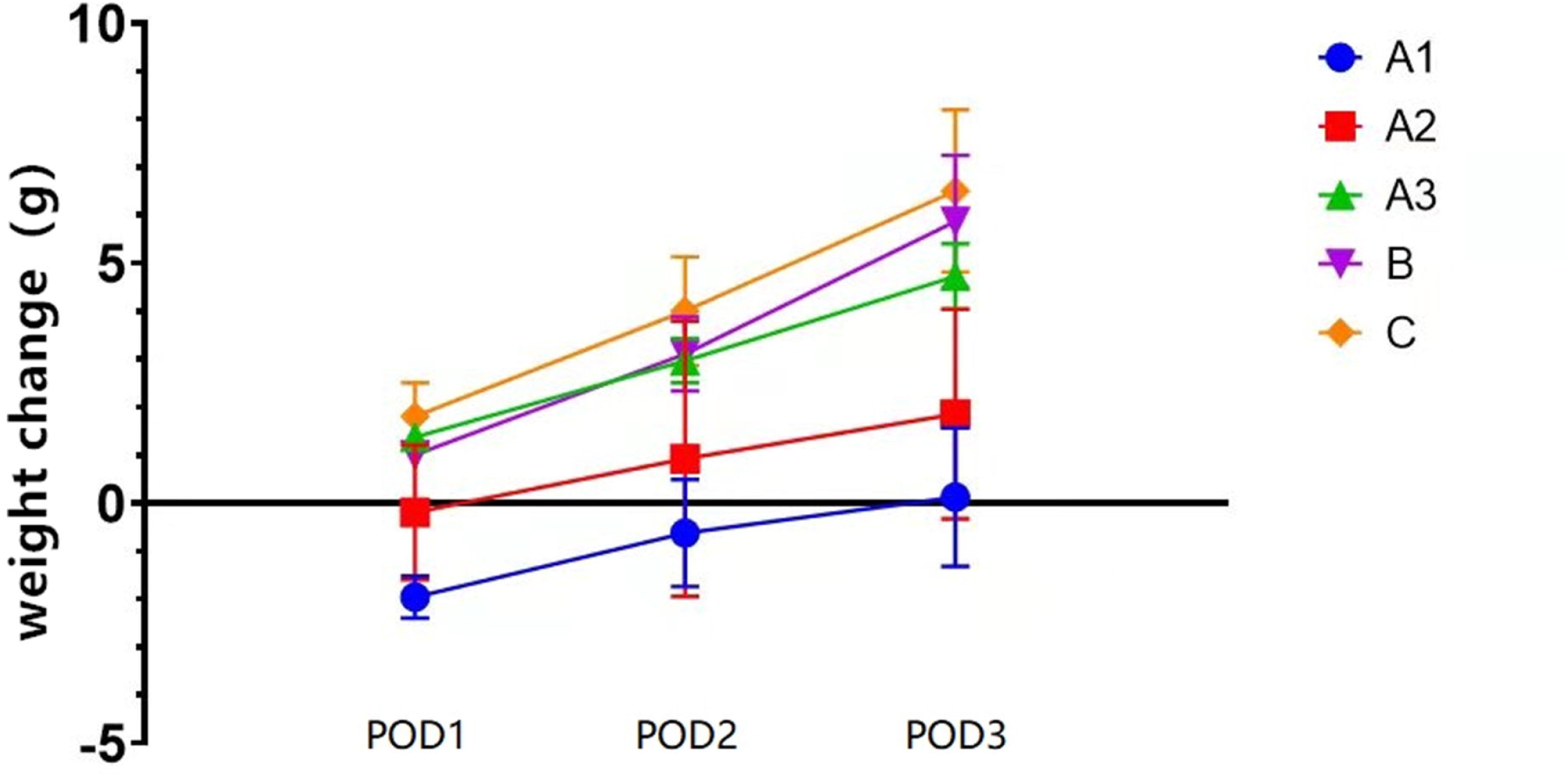
Average increase in weight over 3 days after modeling Weight gain represents the difference between the daily body weight of rats in each group and the body weight before modeling. Body weights were recorded at postoperative day 1 (POD1), POD2 and POD3.

**Figure 4.**
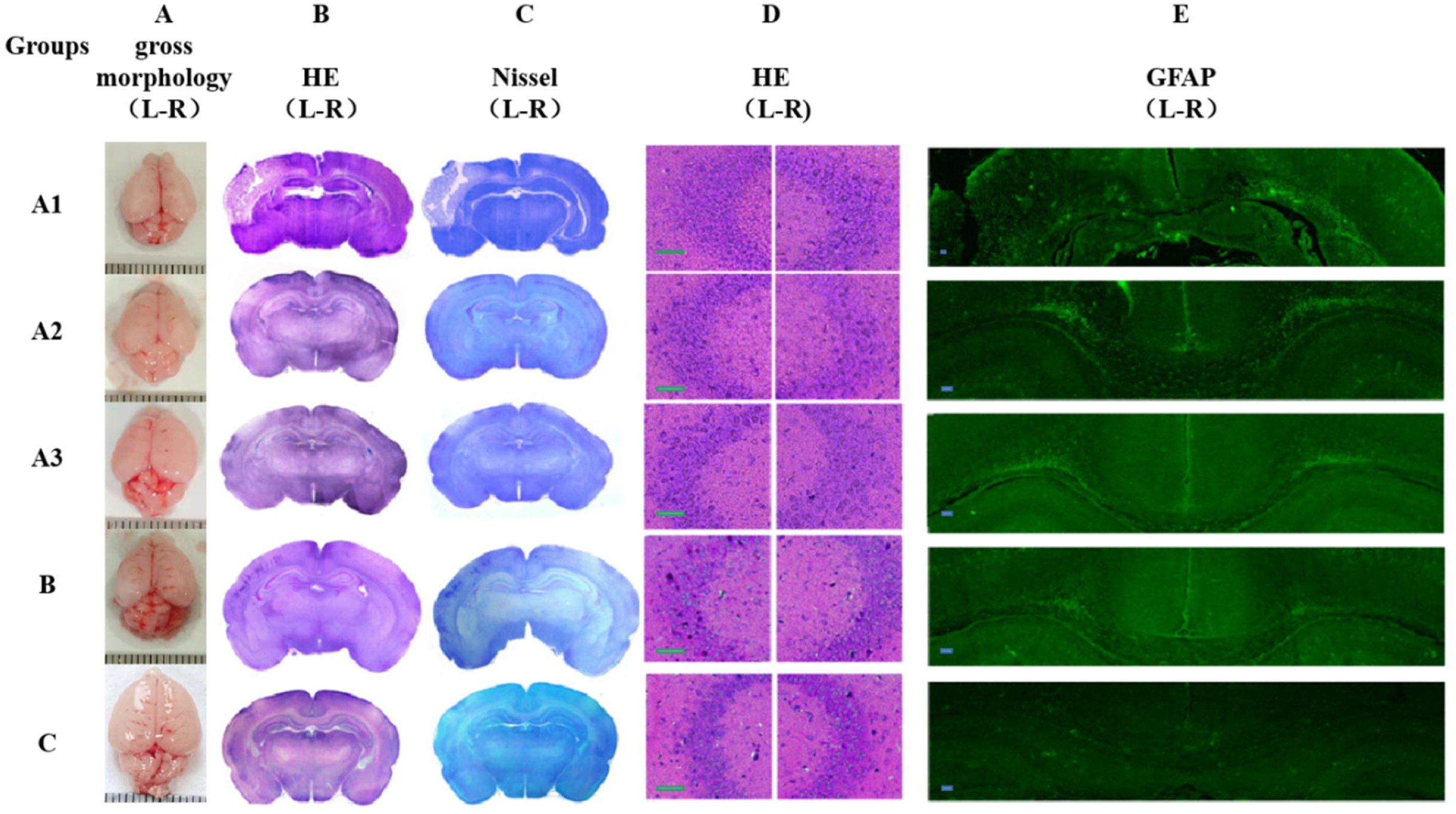
Pathological sections of brain tissue in each group (a) Gross morphology of the brain tissue. The most significant damage was observed in group A1. (b): Brain section (HE-staining) showing significant liquefaction and necrosis in the left brain of group A1 and atrophy of the left brain in group B. (c): Brain section (Nissl staining). (d): Left symmetrical hippocampus (HE-staining), scale bars represent 100μm; (e): Right symmetrical hippocampus (HE-staining), scale bars represent 100μm, Cell count between left and right of the hippocampus was significantly different in group A1,A3 and B. (f): Corpus callosum (GFAP immunofluorescence), scale bars represent 100μm. GFAP positive cells were observed in the corpus callosum of group A2, A3 and B.

### Score of cerebral injury

A1: The liquefaction necrosis area of the left cortex was more severe relative to that of the right cortex (Figure 6). Cells in the left hippocampus were disorganized and the staining of nucleus was light (Figure 7). In addition, the cell count of the left hippocampus was less compared with that of the right hippocampus (p<0.05). Notably, GFAP-positive cells were observed in both left and right brain.

A2: Injuries of left and right cerebral cortex were mild and cells in the hippocampus region were disorganized. Cell count in left and right of the hippocampus was not statistically different (p>0.05) and GFAP-positive cells were observed in the corpus callosum.

A3: The right cortex was atrophic and cells in the right hippocampus region were disorganized. The cell count of the right hippocampus was less compared with that of the left hippocampus (p<0.05). GFAP-positive cells were observed in the corpus callosum.

B: The left brain of this group was severely atrophic. The cell count in the left hippocampus was less relative to that of the right hippocampus (p<0.05). GFAP positive cells were observed in the corpus callosum.

C: No significant changes were observed in the brain tissues of this group. Cells in the hippocampus region were arranged neatly, and no GFAP positive cells were observed in the corpus callosum. Notably, the difference in cell counts between the left and the hippocampus was not statistically significant (p>0.05).

The degree and score of cerebral injury was explored (Table 3). The results showed that group A1 had very severe left hemisphere injury (4.0 points) (Table 6), and mild to moderate right hemisphere injury (1.80 points). This is mainly because the left hemisphere was subjected to permanent ischemia, whereas the right hemisphere was subjected to transient ischemia. Group A2 exhibited mild to moderate left and right hemisphere injury (1.45 points left and 1.36 points right), attributed to induction of transient ischemia on both sides. The duration of hypoxia in group A1 and A2 was 30 minutes and the time to release the bridge was the same (90 minutes). Therefore, these two groups exhibited similar features. Blood flow of the right hemisphere was restored in A1 whereas blood flow was restored in both right and left in group A2. This implies that the degree injury of the left and right hemisphere in A1 was greater compared with that in A2. Group A3 showed mild to moderate right hemisphere injury (1.82 points), and mild left hemisphere injury (1.00 points). Group B exhibited severe left hemisphere injury (3.09 points), and mild right hemisphere injury (1.00 points). The duration of hypoxia in groups A3 and B was the same (120 minutes). The duration of induction of unilateral ischemia in A3 was of 150 minutes whereas rats in group B were subjected to permanent unilateral ischemia. This indicates that unilateral injury was more severe in group B compared with group A3. Notably, if rats are subjected to hypoxia only, the damage would be mild in both B (right) and A3 (left).

**Table 6.** Evaluation of cerebral injury of each group based on the evaluation criteria shown in Table 3.

| degree | score | Group A1<br>(n=10) |  | Group A2<br>(n=11) |  | Group A3<br>(n=11) |  | Group B<br>(n=11) |  | Group C<br>(n=12) |  |
| --- | --- | --- | --- | --- | --- | --- | --- | --- | --- | --- | --- |
|  |  | left | right | left | right | left | right | left | right | left | right |
| Normal | 0 | 0 | 0 | 0 | 0 | 0 | 0 | 0 | 0 | 12 | 12 |
| Mild | 1 | 0 | 2 | 6 | 7 | 11 | 2 | 0 | 11 | 0 | 0 |
| Moderate | 2 | 0 | 8 | 5 | 4 | 0 | 9 | 1 | 0 | 0 | 0 |
| Severe | 3 | 0 | 0 | 0 | 0 | 0 | 0 | 8 | 0 | 0 | 0 |
| Very severe | 4 | 10 | 0 | 0 | 0 | 0 | 0 | 2 | 0 | 0 | 0 |
| Mean score | - | 4.00 | 1.80 | 1.45 | 1.36 | 1.00 | 1.82 | 3.09 | 1.00 | 0 | 0 |

## Discussion

HIE caused by perinatal asphyxia leads to neonatal death. Survivors of HIE present with sequelae of varying severity, including epilepsy, cerebral palsy, motor and cognitive decline, attention deficit, hyperactivity disorder and behavioral disabilities^[12]^. HIE occurs in 1 ~ 8 in every 1000 live births in developed countries and the morbidity is approximately 2.6% in developing countries^[13]^. Currently, no effective prevention and treatment measures are available for HIE. Understanding pathogenesis and development of effective therapy of HIE is thus imperative in improving perinatal medicine.

Reperfusion is a key component of HIE pathophysiology and not merely a continuation of the primary injury. Reperfusion promotes production of large amounts of free radicals and cytokines. Reperfusion injury is thus a major cause of neonatal neurological complications^[10,11,14,15]^.

Studies have presented the concept of “reperfusion” which has two layers in the pathological process of HIE. Compensatory hyper-perfusion of cerebral blood occurs at early stages of hypoxia, resulting in exacerbation of hypoxic condition. It promotes reduction in cerebrovascular autonomic functions followed by decompensated “reperfusion”^[16]^. Further, changing in the lumen of the uterine spiral artery leads to intermittent availability of low placental blood supply in some diseases such as preeclampsia. This repeated low perfusion can result in a low degree of IR injury^[17]^. In addition, IR can occur after rescue treatment of newborns at birth. Therefore, HIE comprises two stages of energy failure. The first stage is primary energy failure, which occurs after hypoxic ischemia whereas the second stage includes calcium overload, free radicals, excitatory amino acids and inflammatory cytokines. The second stage occurs 6-12 hours after birth. The brain undergoes secondary energy failure approximately 48 hours after these pathological processes. Mitochondria undergo oxidative damage. The process is involved in “delayed neurologic injury,” exhibited as apoptosis and necrosis, and may last for weeks^[18–21]^. Therefore, it is necessary to explore the mechanism of IR in HIE pathogenesis.

The Rice-vannucci model is a classical HIBD model and is widely used to simulate the pathological process of HIE^[11]^. However, the model does not mimic reperfusion injury that occurs during HIE. Previous studies reported several hypoxia ischemia reperfusion brain damage (HIRBD) models to explore temporary ischemia. Different methods are used including clamping bilateral CCA, middle cerebral artery occlusion (MCAO) and placental ischemia reperfusion model.

Ashwal et al. used the MCAO technique (suture-occluded method) in P14 rats, using a nylon line passed through CCA to occlude MCA under normoxic conditions to induce IR injury^[22]^. Mitsufuji used electrocoagulation to occlude the right internal and external carotid arteries, and inserted a nylon line into the right CCA to occlude blood supply from the aorta to the cerebrum while clamping the left external and internal carotid arteries to induce brain ischemia for 60 minutes^[23]^. Renolleau conducted electrocoagulation of the left MCA followed by clamping the left CCA for one hour to simulate the pathological process of IR in HIE neonates^[24]^. Derugin established a P7 rat MCAO model of reversible focal cerebral ischemia through the suture-occluded method based on the study by Ashwal et al. ^[25]^. Derugin performed MRI to verify that the blood supply of the middle cerebral artery was almost entirely interrupted after performing the suture-occluded method. Blood flow was partially restored after the thread was removed^[26]^. Zhang et al. established a HIRBD model using miniature arterial clips. Neonatal rats were subjected to hypoxia for 2 hours in a hypoxic chamber with 8% oxygen concentration and the bilateral CCA was temporarily blocked for 30-90 minutes^[27]^. Amara used the suture-occluded method to establish a unilateral, normoxic, focal transient middle cerebral artery occlusion (tMCAO) model using P10 rats^[28]^. Placental ischemia reperfusion model is established by occluding the uterine artery temporarily with an aneurysm clip to simulate hypoxia ischemia reperfusion that mimics the fetal intrauterine status^[29]^. Further, Michelle et al. established a model of delayed cesarean after hysterectomy based on the placental ischemia reperfusion model^[30]^. Placental under perfusion and/or chorioamnionitis associated with pathogen-induced inflammation in early preterm birth was used to develop a model of prenatal transient systemic hypoxia ischemia combined with intra-amniotic lipopolysaccharide to mimic human systemic placental perfusion defects^[31]^. Other MCAO model preparation methods include photochemical method, microembolization method, left cerebral cortical angiectomy, spontaneous intracerebral hemorrhage method, autologous blood injection and collagenase injection^[32]^. Although the MCAO model of rats is universally accepted as a standard model for mimicking focal transient cerebral ischemia, the operation is complicated and challenging to establish insertion diameter and insertion depth. The model can lead to apparent trauma to neonatal rats. Long-term vascular clamping can cause damage to the CCA and affect blood flow. Placental hypoxia-ischemia-reperfusion model has several limitations including difficult operation, complex process, and it can easily cause preterm birth, miscarriage and stillbirth in rats, as well as loss of fertility in female rats.

Compared to these methods mentioned above, CCA muscle bridge operation does not result in injury of vessels. This method is a straightforward, easy to learn and highly reproducible process. Therefore, we believe this method is an effective HIBD model. Hemodynamics analysis using LSI showed that the CCA muscle bridge is reliable and stable. The results based on A1-1 group showed that the muscle bridge method was similar the Rice-Vannucci method in effectiveness of blood flow blockage. Results for group A2-1 and A2-2 were used to verify stability of the bilateral muscle bridge operation and reliability in establishing whole brain reperfusion injury models. Findings on group A3-2 indicated that blood flow of reperfusion was restored to the preoperative level after the removal of muscle bridge. However, keeping muscle bridge for a long time can affect the status of CCA. Removal of the muscle bridge after 24 hours showed formation of a clot in the CCA resulting in vascular blockage (data not shown). Removal of the muscle bridge within 4 hours can prevent thrombosis despite the possibility of vascular stenosis in our preliminary tests (data not shown). These results show that the muscle bridge method is effective for blood flow blockage. Notably, blood flow of reperfusion is not affected if the muscle bridge is removed after a certain period of time.

Group A2, A3 and B showed similar mortality with a slower body weight gain. Group A1 exhibited the highest mortality as well as negative body weight gain owing to severe brain damage. In our study, the brain tissues were examined on the third day to evaluate the extent of injury through pathological analysis. Infraction volume increases with increase in reperfusion time after the modeling of middle cerebral artery occlusion, and the injury reaches a peak on day 3 after modeling^[33]^. HE and Nissl staining were used to explore moderate and severe brain damage. GFAP immunofluorescence is a more sensitive method for mild brain damage. The earliest and most significant features of hypoxia ischemia injury include change in astrocytes. GFAP is a specific marker protein of mature astrocyte, thus it can be used as an indicator of mild injury. Therefore, brain damages were evaluated by analysis of gross morphology and pathological sections of brain tissues (HE-staining, Nissl staining and GFAP immunofluorescence). Brain damages are classified as normal, mild, moderate, severe and very severe based on these indicators. The findings showed that the left-brain injury was very severe in group A1, which was consistent with the results on body weight. This can be attributed to the interruption of contralateral blood supply in the left cerebral by the right muscle bridge. Therefore, A1 group can be used for comparative study of ischemic and reperfusion brain injury. Group A2 showed bilateral brain damages, thus it can be used as a reperfusion bilateral cerebral injury model. Group A3 represented a unilateral cerebral IR injury model, which can be used for comparison with the HIBD model established using the Rice-Vannucci method. These findings indicate that an appropriate model should be selected for study of HIBRD.

In summary, the findings of our study present effectiveness and stability of the proposed model. The CCA muscle bridge operation is a reliable and stable method for establishing HIBRD model for the study of IR in HIE.

## Author contributions

Xiaoqin Fu, Tianlei Zhang and Wei Lin anddrafted and revised the article and should be considered co-first authors. All authors report no conflicts of interest in this work.

